# RSQ: a statistical method for quantification of isoform-specific structurome using transcriptome-wide structural profiling data

**DOI:** 10.1101/043232

**Authors:** Yunfei Wang, Xiaopeng Zhu, Ming Sun, Yong Chen, Yiwen Chen, Shikui Tu, Boyang Bai, Min Chen, Qi Dai, Haozhe Wang, Michael Q. Zhang, Zhenyu Xuan

## Abstract

The structure of RNA, which is considered to be a second layer of information alongside the genetic code, provides fundamental insights into the cellular function of both coding and non-coding RNAs. Several high-throughput technologies have been developed to profile transcriptome-wide RNA structures, i.e., the structurome. However, it is challenging to interpret the profiling data because the observed data represent an average over different RNA conformations and isoforms with different abundance. To address this challenge, we developed an RNA structurome quantification method (RSQ) to statistically model the distribution of reads over both isoforms and RNA conformations, and thus provide accurate quantification of the isoform-specific structurome. The quantified RNA structurome enables the comparison of isoform-specific conformations between different conditions, the exploration of RNA conformation variation affected by single nucleotide polymorphism (SNP),and the measurement of RNA accessibility for binding of either small RNAs in RNAi-based assays or RNA binding protein in transcriptional regulation. The model used in our method sheds new light on the potential impact of the RNA structurome on gene regulation.

## Background

RNA carries regulatory information not only within its primary sequence,but also within its secondary structures [1, 2].The RNA secondary structure is fundamental to RNA transcription, splicing, localization and turnover [1, 3–9], and several profiling methods have been developed [10–20], mostly based on the technologies of applying chemical or enzymatic probes to identify the states of a bases of RNA as either single-strand, doublestrand or solvent-exposed [21, 22]. The coupling of next-generation sequencing technology to these methods has allowed them to be adapted to the scale of the whole transcriptome, yielding the first glimpse of the ‘RNA structurome’ [10–19], and raising the question of how structured regions control RNA functions and gene expression [2].

Several computational methods have been developed to reconstruct the structurome from RNA structural profiling data. For example, the SeqFold method uses the Boltzmann sampling method to generate a pool of RNA conformations,and clusters them into groups.The cluster centroid nearest to the structural profiling data is then considered as the structure of a gene [23]. MaxExpect integrates the free energy model with constraints inferred from RNA structural profiling data to predict structures with maximal expectation [24]. The RNAStructure method is designed for SHAPE data, which uses RNA structural profiling data as prior knowledge to constrain the RNA folding procedure [25].These methods show good performance when used to predict the optimal structure given a single RNA sequence with experimentally inferred constraints.

However, the circumstance on which these methods are based only occurs when the gene is transcribed into a single transcript. In eukaryotes, it is very common that many genes can produce multiple isoforms of transcripts through alternative splicing [26] and alternative promoters [27, 28]. For example, nearly 80% of protein-coding genes have multiple isoforms(GENCODE V19 annotations [29]). For those genes, the RNA structural signals captured in the profiling analysis result from a mixture of structures folded by all expressed isoforms with potentially different levels of abundance. In addition, around 5% of protein-coding genes and 2.8% long intervening non-coding RNAs (lincRNAs) overlap with other genes in the exonic regions. The complexity of transcriptome makes it difficult to accurately quantification of RNA structures due to potential ambiguity of mapping the short sequencing reads from structural profiling data to the original isoform. (Figure 1A).

Additional challenge in RNA structural profiling data analysis could come from the discovery that a single RNA sequence may fold into multiple conformations. In prokaryotes, multiple conformations are shown to be involved in the regulation of translation initiation [30] and protein synthesis [31, 32].In eukaryotes, different conformations of a RNA mediated by RNA-binding proteins (RBPs) can change the accessibility of the RNA for other small regulatory RNAs, and switch the modes of gene translation [33, 34]. RNA conformation variation is also evident in high-throughput RNA profiling data, in which some RNA bases at certain positions show strong conflicts between single-strand and double-strand signals (Figure 1B). Some methods, such as Sfold [23], CENTROIDFOLD [35] and SeqFold [36], considered this conformation variations in their model and achieve improved prediction performance.

This has motivated us to develop RNA structurome quantification (RSQ), a method based on a statistical model to systematically integrate transcriptome-wide RNA structural profiling information into RNA structurome modeling and quantification, while considering both the alternative isoforms and conformations(Figure 1C). RSQ can analyze data from all mainstream high-throughput RNA structural profiling technologies, including PARS, FragSeq, SHAPE-Seq, DMS-Seq, icSHAPE, and also conventional low-throughput structural profiling data. We found that RSQ can interpret RNA structural profiling data better than the other existing method. The quantified RNA structurome can reveal the diverse roles of RNA structures in translation efficiency as well as in transcription initiation accuracy. RSQ also provides useful information for measuring RNA accessibility,which is essential for the identification of RBP targets, accurate interpretation of endogenous miRNA regulation, and rational designs of small interfering RNAs (siRNAs),antisense oligonucleotides and trans-cleaving ribozymes in gene knockdown studies.

**Figure 1.**
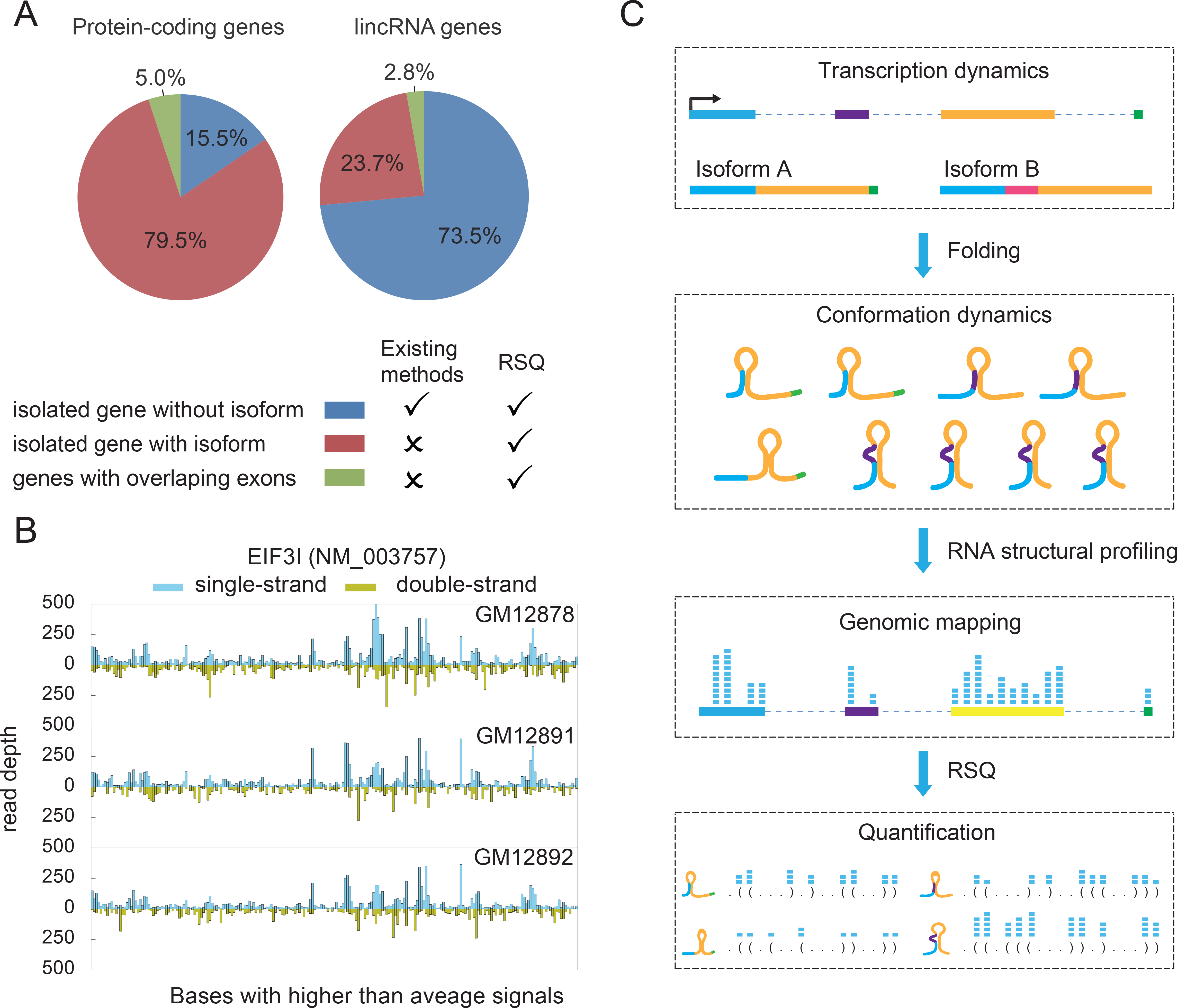
Two layers of dynamics in the RNA structurome and schematic model of the RSQ method.(A)Gene transcription dynamics for protein-coding genes and lincRNAs.(B)RNA conformation dynamics. Taking EIF3I as an example, RNA structural profiling data show strong conflicting signals from single-strand and double-strand data in some regions,which are common among cell lines in a family trio.(C)Schematic workflow of RSQ method. RNA structural profiling technologies capture signals from a mixed pool of RNA conformations that are folded from all expressed isoforms for a given gene,losing the information of the conformation from which they are captured. To recover the information, the reads are piled onto the genome, and the EM algorithm is used to reassign the reads to the RNA conformation pools with the maximum probability

## Methods

### Generative model for single strand structural data using Expectation Maximization (EM) algorithm

The least complicated case is a gene only transcribes a single transcript. We assume that the transcript with length *L* can fold into *K* conformations, and a set of reads from RNA structural profiling data (denoted as R)are uniformly and independently sampled from all positions in the single strand structures of the transcript, in total *N* reads. Then, similar to the RNA-Seq isoform quantification problem described in published studies [37, 38], an extended generative model for the EM algorithm can be constructed to assign the *N* reads to the *K* conformations. Each of the *N* reads can be associated with a latent variable *Z_d_* = *j*, *j* ∈ {1,…, *K*}, indicating that the *d*th read is derived from conformation *j*. The primary parameters of the model are ***θ*** = (*θ*_1_…,*θ_K_*), which correspond to the percentages of reads derived from every conformation. Given the conformation assignment ***Z***, the likelihood of observing all reads ***R*** with parameter ***θ*** can be written as

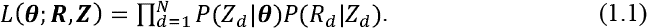

Under the uniform distribution assumption, the *N* reads can be grouped according to their coordinates on the transcript. We define a vector ***X***={*X_1_, X_2_,…, X_i_*, …, *X_L_*} to represent the number of reads mapped at each position of transcript, and we have 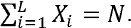 Similarly, the latent variable *Z_d_* for read *R_d_* mapped to the same position can be reshaped to an indicator matrix *Z_ij_,* where *Z_ij_* = 1 if *R_d_* is mapped at position *i*, and *R_d_* is derived from conformation *j(Z_d_=j).* A conformation profile matrix is also defined as *I_ij_*, where *I_ij_*=1 denotes that the *i*th position of the *j*th conformation is in a single-strand structure; otherwise *I_ij_*=0. To simplify the denotation, let *L_j_* denotes the total length of singlestrand structure (called effective length) of the *j*th conformation, where 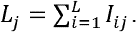

Then the conditional probability of *Z_ij_*=1 given ***θ*** is

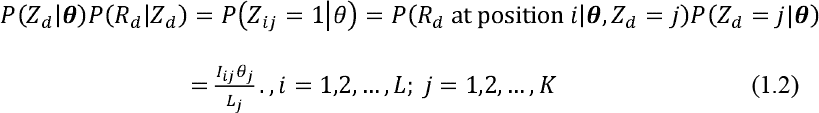

### Inference with the EM algorithm

With the conformations and RNA structural profiling data for a transcript, the aim is to infer the model parameters ***θ*** or alternative parameters **π**=(*π*_1_,…,π_***K***_), which correspond to the relative expression level of the *K* conformations. It is equivalent to inferring either ***θ*** or **π**. Whichever one is easier to be estimated, the other one can be calculated according to the relationship described below:

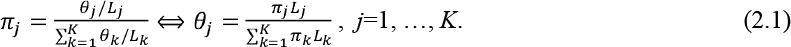

Here, we choose to infer the parameters ***θ***, which can be estimated by maximizing the likelihood of the observed data:

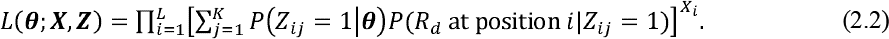

We use the EM algorithm to find the maximum likelihood values for ***θ***. In the expectation step, the expected value of the log-likelihood function is calculated with respect to the conditional distribution of latent variable ***Z*** given *X* under the current estimate of the parameters ***θ***. In this case, the expected values of *Z_ij_*, denoted as *Z_ij_*, can be calculated by

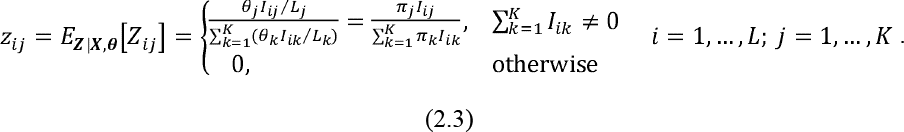

The expected value of the log-likelihood can be written as

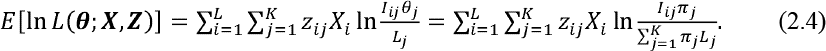

In the maximization step, equation (2.4) is maximized with respect to *π*^(*t*)^, together with the constraints that 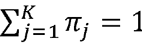:

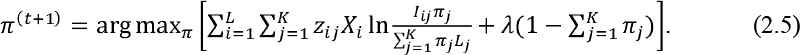

Solving the above equation yields the updated parameters (Supplementary document S2.1):

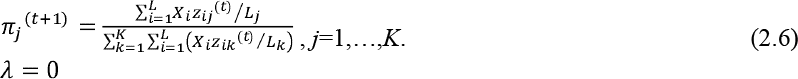

The iteration runs until it reaches convergence 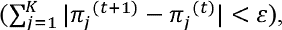 or exceed the preset maximum number of iterations. Because the parameters in *p*(*R_d_*)at position *i*|*Z_ij_* =1 are predetermined by the conformation profiles, and thus the observed data likelihood is concave, the EM algorithm is guaranteed to find the optimal value *Z_ij_* * to maximize likelihood [38].

When the EM algorithm reaches convergence, the term

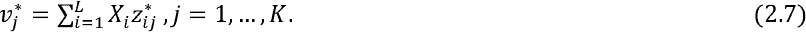

is actually the total number of reads assigned to the *j*th conformation. From equations (2.1), the relative read abundance *θ_j_* can be written as

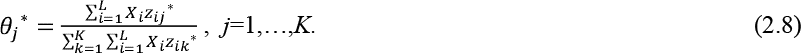

Then the abundance of the *j*th conformation can be estimated by

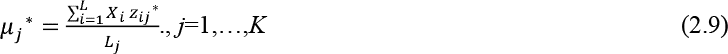

### Model generalization: genes with multiple isoforms

Genes with multiple isoforms require additional processing before applying the EM algorithm. Given a gene with *M* isoforms and total *K* conformations, we introduce the EFastS format (Supplementary document 1) in order to apply the EM algorithm to the conformations of these isoforms. The exons from *M* isoforms are collapsed to obtain a gene model which transcribes a single union transcript with length of *L*. For each of *K* conformation, if the position in the union transcript is excluded, it will be substituted with hyphen. (Figure 2A). The gene model and *K* aligned conformations can be used as the input for the generative model described above. When the EM algorithm converges, the expression level of the *l*th isoform, *E_l_*, can be directly estimated from all the conformations belonging to this isoform (*C_l_*),.

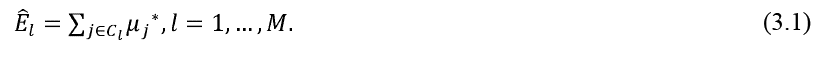

Besides directly estimating *E_l_*, the expression level of an isoform of a gene can be independently obtained from other measurement, such as transcripts per million reads (TPM) from RNA-seq. In that case, we could calculate the relative expression level of the *l*th isoform P*_l_* by

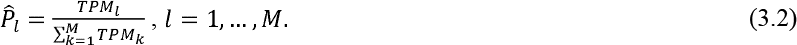

and use it as constraints for parameter **π** in our EM algorithm by

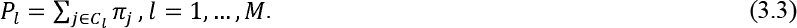

Taking equation (3.3) as the Lagrange multiplier in the maximization step yields:

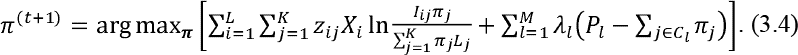

The updated parameters (details in Supplementary document S2.2) are

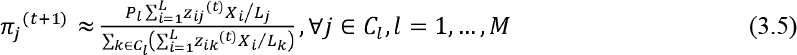

**Figure 2.**
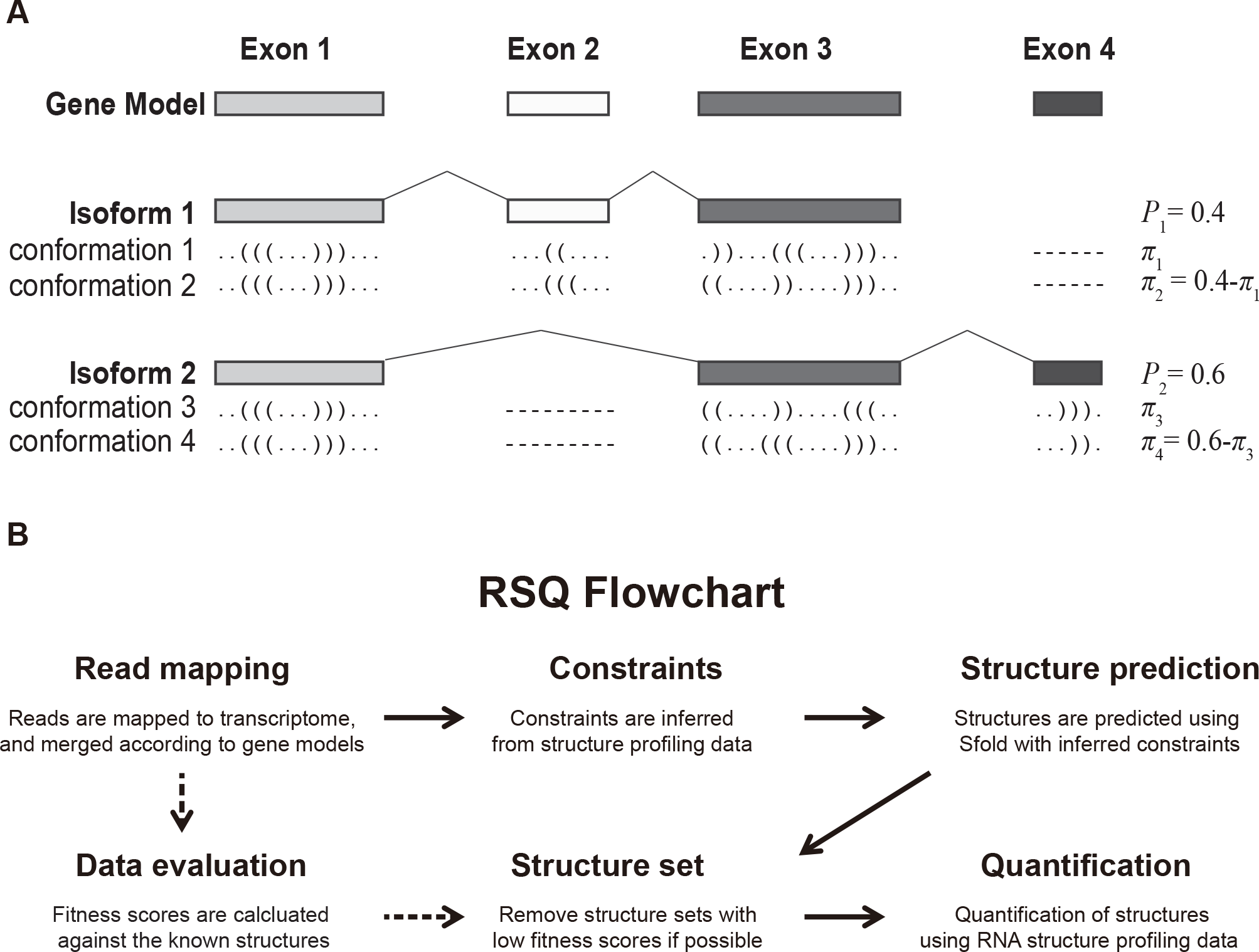
RSQ method. (A) Collapsing genes with multiple isoforms. P*l* is the relative transcript abundance for the *l*th isoform and *πj* is the percentage of *j*th conformation. (B) Flowchart of RSQ method.

### Model generalization: genes with both single-strand and double-strand information

For RNA structural profiling technologies producing both single-strand and doublestrand information, such as PARS data, the likelihood function can be written as the product of the likelihood functions of both the single-strand and double-strand data. Let *S* and *D* denote the sets of reads, and ***X_S_***= {*X_S,i_, i=1,…,L*} and ***X_D_***={*X_D,i_,i*=1,…,*L*} and as the number of reads mapped at each position from single-strand and double-strand data, respectively. The complete log-likelihood function is

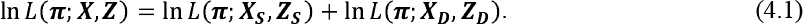

Here, ***Z_S_*** and ***Z_D_*** are indicator matrices for reads in *S* and *D* sets separately. If genes have multiple isoforms, the constraints in equation (3.3) hold for both data types, given the estimated *P_l_* from RNA-Seq data. Finally, maximization of the expected value of the complete log-likelihood function can be represented as

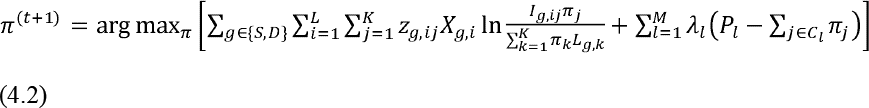

The profile matrices (*I_S,ij_, I_D,ij_*) and effective length (*L_S,j_, L_D,j_*) are for single-strand and double-strand structures in *j*th conformation respectively. The updated parameters are approximated (details in Supplementary document S2.3) as

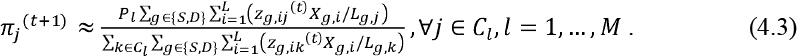

It should be noted that the total number of reads for each sample should not be scaled to the same because theoretically the samples with more observations (or reads) are supposed to contribute more during parameter estimation. Besides, reads for singlestrand and double-strand data may not contribute equally in the model, so weight can be applied to each sample before implementing the EM algorithm. The weights can be either obtained from prior knowledge, such as characteristics of RNase S1 and V1, or empirically estimated from the current data, such as fitness scores (Supplementary document 3.2) or predictability to known structures. For example, when the average as fitness scores 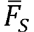 and 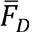 to known structures are used to weight two samples for single and double-strand information respectively, the updated parameters can be written as

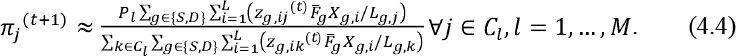

### Model generalization: Combining replicates from single and double strand data

When RNA structural profiling data have biological replicates, the likelihood function can be written as a product of the likelihood functions of all the replicates. When weights are available, the reads can be adjusted by weights and then combined to run RSQ algorithm. Let S’ and D’ denotes the weights adjusted reads for single-and double-strand data, respectively, the complete log-likelihood function then can be written as

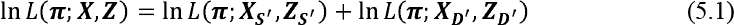

The updated parameters are similar to (4.3)

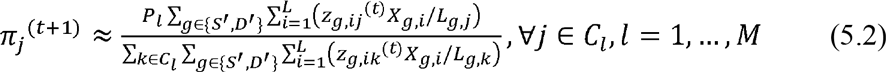

### Simulation

Theoretically, the sequencing reads were generated for a transcript or isoform from a mixture of conformations according to the relative conformation abundance π. To simulate this process, 100 RNAs were selected from the *S. cerevisiae* transcriptome, and conformations were predicted using Sfold [39]. For each RNA, the parameters ***π*** was preset and the relative read abundance ***θ*** was calculated based on (2.1). To evaluate how sequencing depth in structural profiling could affect the quantification performance, varied number of reads were assigned to the conformations according to ***θ***, and then randomly distributed to all effective positions of each conformation. Finally the RSQ method was applied to estimate 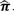. The average difference between the preset and estimated parameters was calculated to evaluate the performance of RSQ. To simulate the noise effects, varied percentages of noisy reads were randomly assigned to all the positions of a given transcript. When combining single-strand and double-strand reads, the two types of reads were sampled independently and combined to run RSQ. For the genes with multiple isoforms, the relative isoform expression level *P_l_* was calculated by taking the sum of the preset *π_j_*,∀ *j* ϵ *C_l_*.

### RNA accessibility

Based on the quantification results of RSQ, the accessibility of a given transcript can be quantified. Let *A_ij_*=1 denote that the *i*th position of the *j*th conformation is accessible; otherwise, *A_ij_*=0. If transcript *T* has a miRNA binding target located from *b_s_* to *b_e_*, 1≤*b_s_*<*b_e_*≤*L*, then its overall accessibility for the miRNA is calculated by

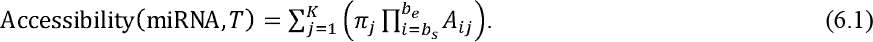

The product of *A_ij_* from *b_s_* to *b_e_* means that a continuous region is considered to be accessible only when all the positions within are accessible.

### Implementation

RSQ was implemented as a Python package with C/C++ modules for RNA structural profiling data analysis. It is publicly available from Python Package Index https://pvpi.pvthon.org/pvpi/rsq. RSQ provides a general solution from the raw RNA structural profiling data to the quantified RNA structurome (Figure 2B). RSQ has defined universal data formats, such as FastD, FastC, FastS and EFastS (see Supplementary document 1). These data formats are compatible with mainstream RNA structural profiling technologies. Documentation for data input, parameters and output are provided for the users of RSQ to easily generate the input files, tune the parameters according to their requirements and interpret the output results. Demonstration examples are also provided with the RSQ package.

## Results

### Evaluation of RSQ method

Simulation data were used to evaluate the performance of the RSQ method. 100 transcripts were selected from the *S. cerevisiae* transcriptome, and structures were predicted using Sfold [39]. The simulation results showed that RSQ exhibited excellent performance in deciphering the structural dynamics, even for genes with very low coverage (Figure 3A). RSQ also showed good tolerance to noise. Given data with noise levels up to 30%, more than 75% of predictions showed less than 10% difference when compared to the preset percentages (Figure 3B). In addition, the sample balance effect was evaluated by varying the ratio of the amount of single-strand and double-strand data. Although samples with balanced read coverages for single-and double-strand data showed slightly better performance when we assume the weights of both types of data are equal, the proportion of single-strand data doesn’t affect the overall prediction performance significantly. (Figure 3C). With the same amount of reads, RSQ was performed better for genes with single transcript than those with multiple isoforms. When applying the independently measured expression levels of isoforms as constraints, the performance of RSQ can be improved. (Figure 3D).

**Figure 3.**
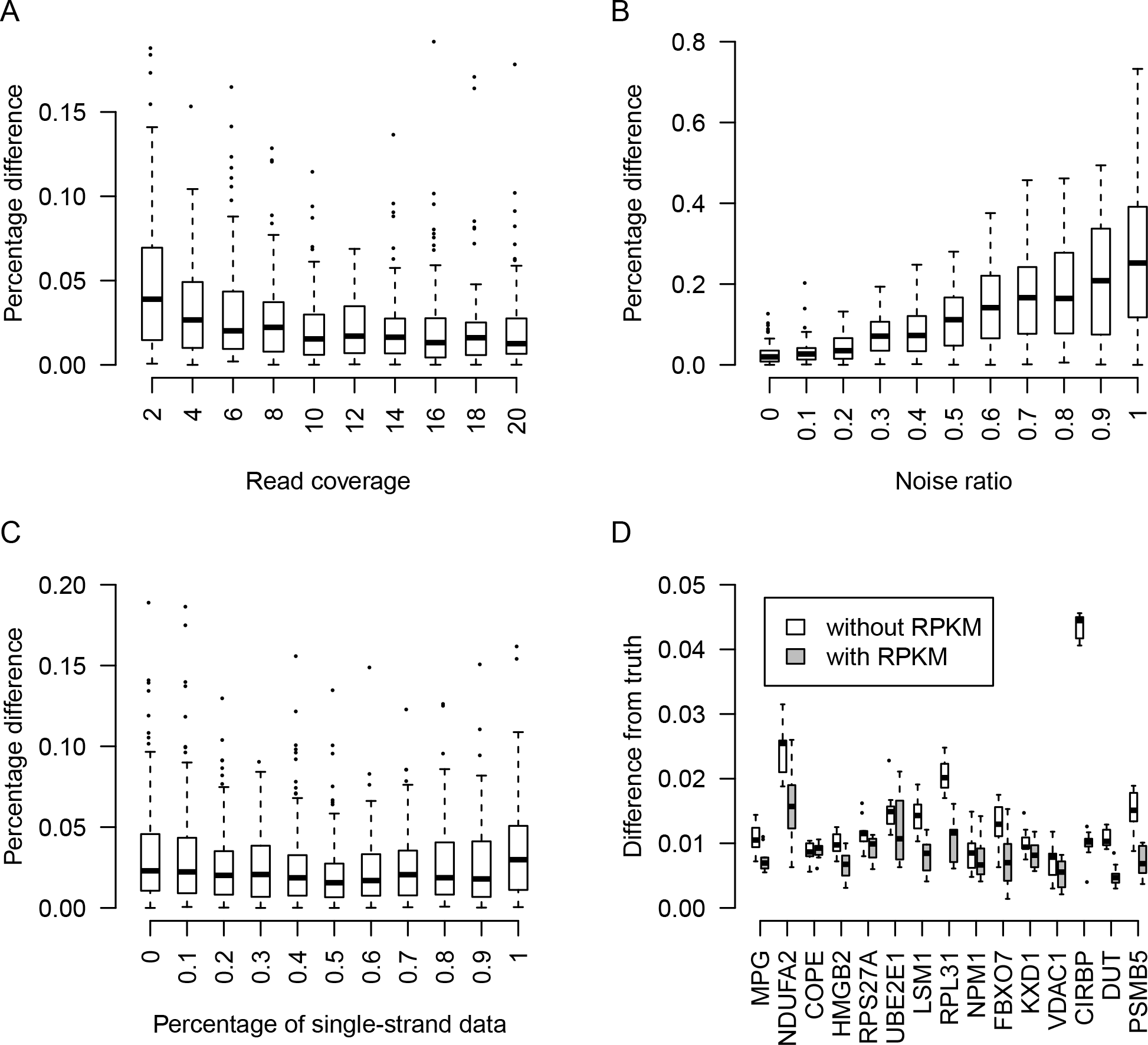
Boxplots of RSQ performance based on simulated data. RNA structures were predicted by Sfold [45] for 100 S. cerevisiae RNAs. (A) Read coverage effects for RSQ performance. Given RNA structures, read coverage ranging from 2 to 20 was randomly simulated from singlestrand structures. (B) Noise tolerance of RSQ method. The read coverage was set to 10. Varied percentages of noisy reads were generated randomly along the RNA sequence (100% means the amount of noise reads is equal to the amount of structural reads). (C) Single-strand and doublestrand data balance effect of RSQ method. The percentage of single-strand reads ranged from 0(double-strand reads only) to 1 (single-strand reads only). The total read coverage were 10 and no noise was added. (D) Performance for 15 genes with multiple isoforms. With isoform expression levels (TPM) used as constraints, the performance of RSQ was improved.

### Profiling efficiency evaluation of existing RNA structural profiling data

To assess profiling efficiency of a variety of RNA structural profiling approaches [10, 11, 16, 18, 19], fitness scores were calculated in yeast cell lines and in several human cell lines to evaluate the fitness of the RNA structural profiling data to the known structures (Figure 4). For the *in vitro* PARS data with both single-strand and double-strand information, the single-strand data always fit better than the double-strand data, which might result from the lower accessibility of RNase to double-strand regions. Although PARS data had additional double-strand information, when fit to known structures, the PARS data fit worse than *in vivo* DMS-Seq data. This indicates that the structural information captured *in vitro* was less representative of the real structural profiles.

**Figure 4.**
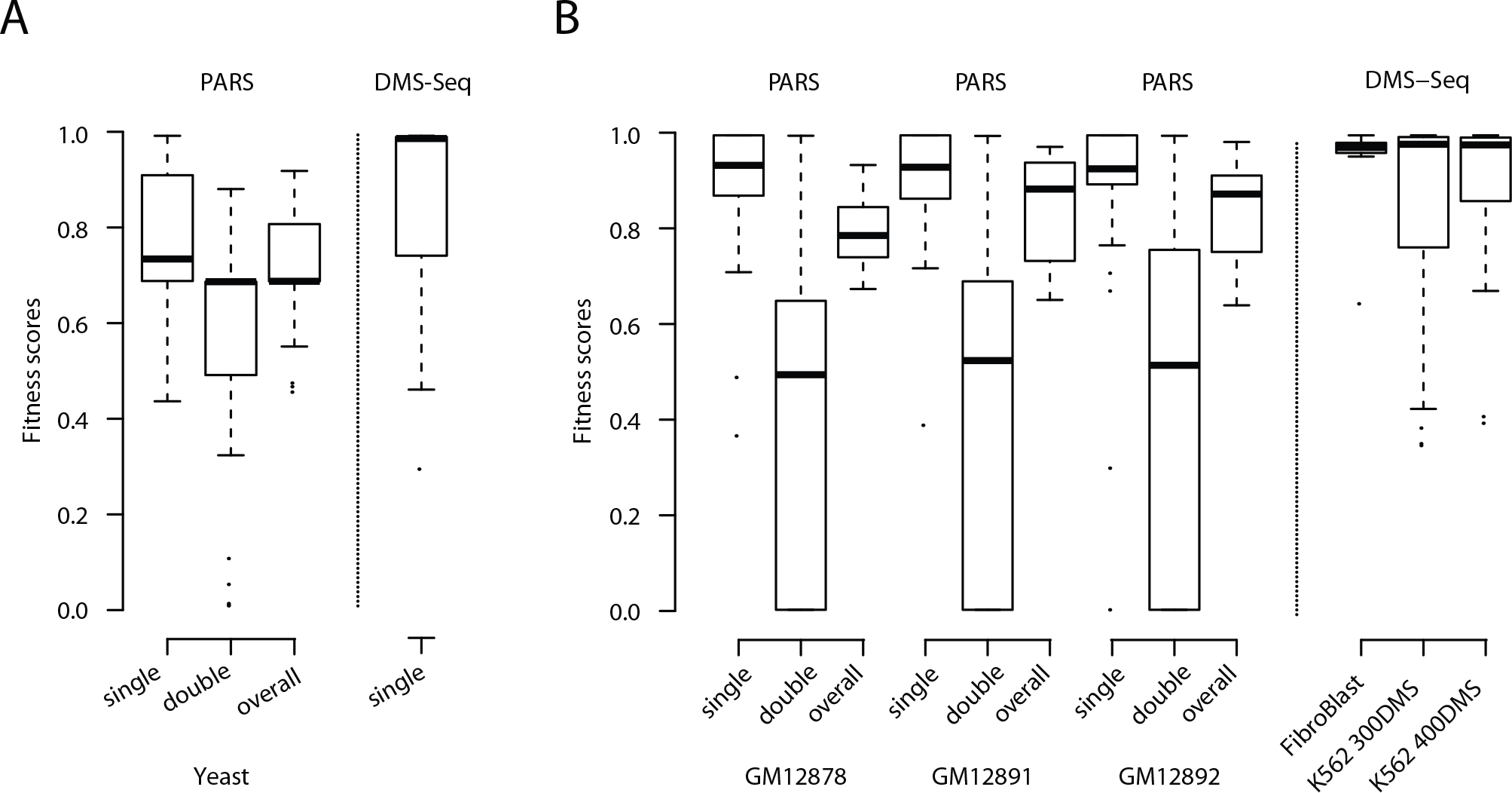
Fitness analysis of existing RNA structural profiling data with respect to known RNA structures. PARS produced both single-strand and double-strand data, and DMS-Seq has only single-strand information. (A) Fitness score distribution of RNA structural profiling data in yeast.(B) Fitness score distribution of RNA structural profiling data in human cell lines. For DMS-Seq data in K562 cell line, DMS was applied in two different concentrations.

### RNA structurome profile in human cell lines

The human PARS data [10] for a family trio and DMS-Seq data [11] in fibroblast and K562 cell lines were used to explore the RNA structurome profile among cell lines. The data were analyzed by RSQ using the default parameters. The RNA conformation cluster profiles inferred from experimental data are different from those inferred from theoretical predictions, apparently because some of the theoretical structural clusters are not favored in physiological conditions. Additionally, without the context of *in vivo* interactions, such as RBPs and small RNA interference, the theoretical minimum free energy (MFE) structure may not be the optimal structure in physiological conditions (Figure 5A). Certain clusters of the theoretical conformations are less observed in physiological conditions. Moreover, for each gene, theoretical structures that were not favored in one cell line might be favored in some other cell lines, indicating that the structurome also varies among cell lines in the conformation compositions (Figure 5B).

The RSQ quantification result for the family trio showed that the percentages among cell lines correlated well, even between samples without a blood relationship (Figure 5C). Consistent with the structural percentages, the structural expression levels estimated from either single-strand or double-strand information also correlated well among cell lines (Figure 5C). These facts indicate that RNA conformation composition is a relatively stable feature in the RNA structurome among cell lines. To determine whether the variation in RNA conformations resulted from the variation in gene expression levels among cell lines, the relationship between gene expression abundance and RNA conformation composition was assessed (Figure 5D). The results showed that the difference in the RNA conformation composition does not directly result from the gene expression level variation among cell lines, indicating that other cell-specific regulatory events, including RBPs and small RNA interference, were involved in the determination of the RNA conformation composition.

**Figure 5.**
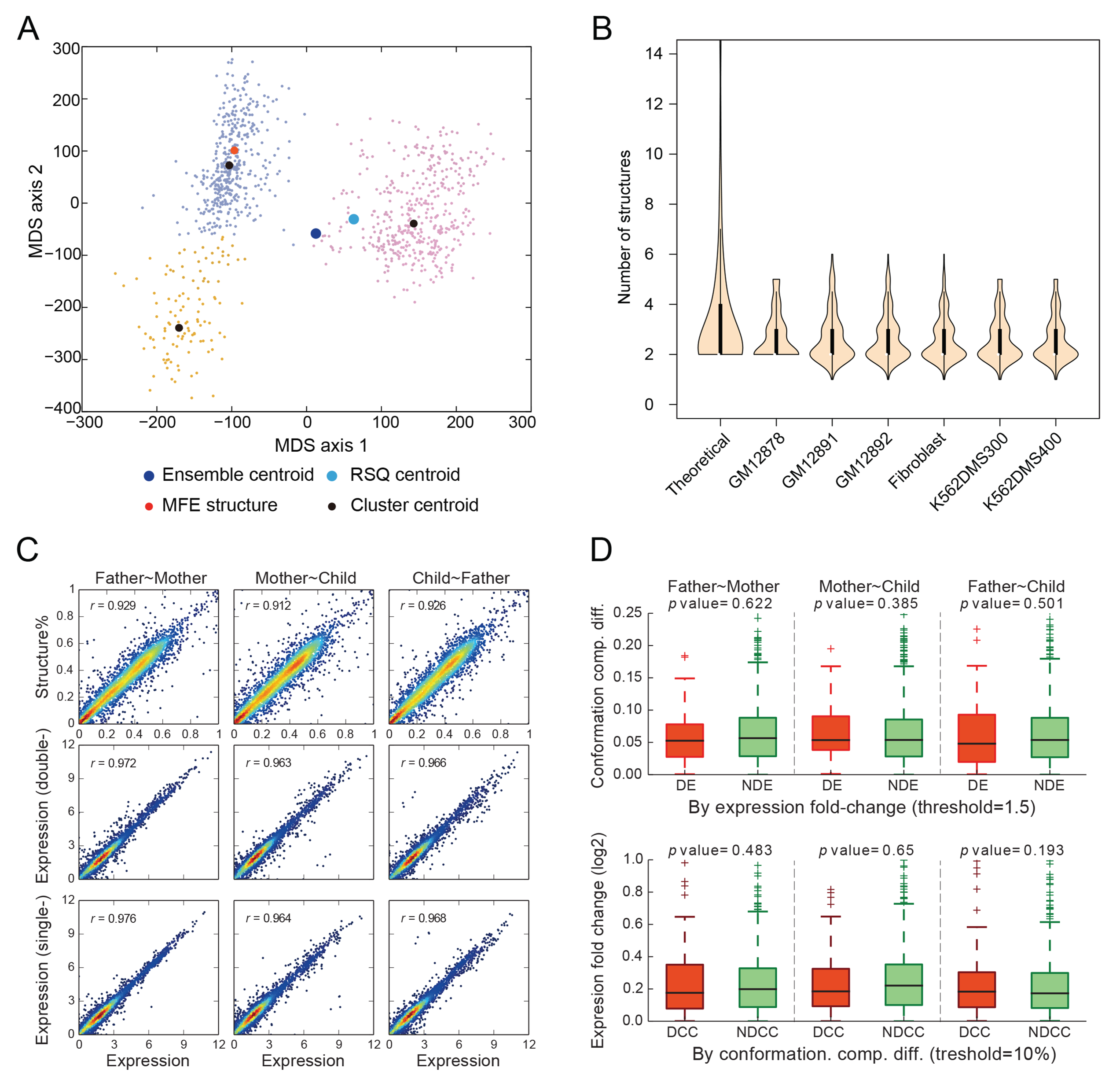
Characteristics of RNA conformation dynamics in human cells. (A) Multidimensional scaling (MDS) of the clustering for 1,000 structures. The theoretical structures were grouped into 3 clusters. By applying RSQ to RNA structural profiling data, only two clusters were supported by experimental data, and the minimum free energy (MFE) structure is not favored in experimental conditions. (B) Violin plot of number of theoretically predicted and experimentally supported clusters. Non-isoform genes with length < 3kb and coverage> 5nt in all samples were Wang Page 28 Quantification of RNA structurome using next generation RNA structural profiling data used. (C) Scatterplot of structural percentages and gene expression levels among a family trio. Pearson correlation coefficient was shown. (D) Relationship between differential expression and differential conformation composition. Genes are split into two groups by whether they are differentially expressed (DE, absolute fold-change>1.5) or not (NDE, the rest) between samples, and then the RNA conformation composition difference (defined as maximum absolute difference of structural percentage for a given gene) was calculated for the two groups. The results show that the RNA conformation composition difference is not significant (Student’s ttest) between DE and NDE genes (upper panel). In parallel, the genes are also split into two groups by whether they have different conformation composition (DCC, maximum absolute structure percentage difference >10%) or not (NDCC, the rest), and the absolute values of gene expression fold change (log2) are calculated for both groups. Similarly, no significant difference in expression fold change (Student’s t–test) is observed between the DCC and NDCC groups (lowerpanel).

### RNA accessibility

Using the RSQ method, RNA accessibility for small interfering RNAs and RBPs can be quantitatively assessed in single base pair resolution. Taking the PARS data in the GM12878 cell line as an example, RNA accessibility was calculated for 100 genes that only have single transcript. For comparison, RNA accessibility was also calculated for the top structures to mimic the SeqFold method [36] (Figure 6A). In general, the region right after the transcription start site (TSS) showed much higher accessibility, which reflects the prevailing occupancy of the Polymerase II transcription complex at the promoter region [36]. In addition, the top structures only contributed around half of the overall accessibility on average, indicating that RNA accessibility was inappropriately estimated (the top structures were exaggerated while the other structures were ignored) when only the top structure was considered for each transcript.

To evaluate the effect of RNA conformation dynamics to miRNA-mRNA interaction, the accessibility scores were calculated for miRanda [40] predicted miRNA targets in the human transcriptome [41]. The results showed that although these predicted miRNA binding events have good scores when using the mirSVR algorithm [42] and have been conserved during evolution, only a portion of the binding sites were accessible in individual cases (Figure 6B).

Constrained by the unique conformation assumption, some of the RBPs and small RNA interference events might be neglected. For example, hsa-miR-302a has a miRanda predicted binding site at the 3’ UTR of the COX7B gene, but the binding site is only accessible in the secondary optimal conformation, which contributes 31.7% of all the COX7B transcripts (Figure 6C). This interaction would be neglected if the accessibility of the optimal structure were used to represent that for all the COX7B transcripts. In addition, the quantified RNA conformation dynamics assist in interpreting small RNA regulation efficiency. In the case of COX7B gene, as most (68.3%) of its RNA molecules are not accessible at the has-miR-302a target site, the miRNA regulation efficiency is greatly reduced in the corresponding cell line.

**Figure 6.**
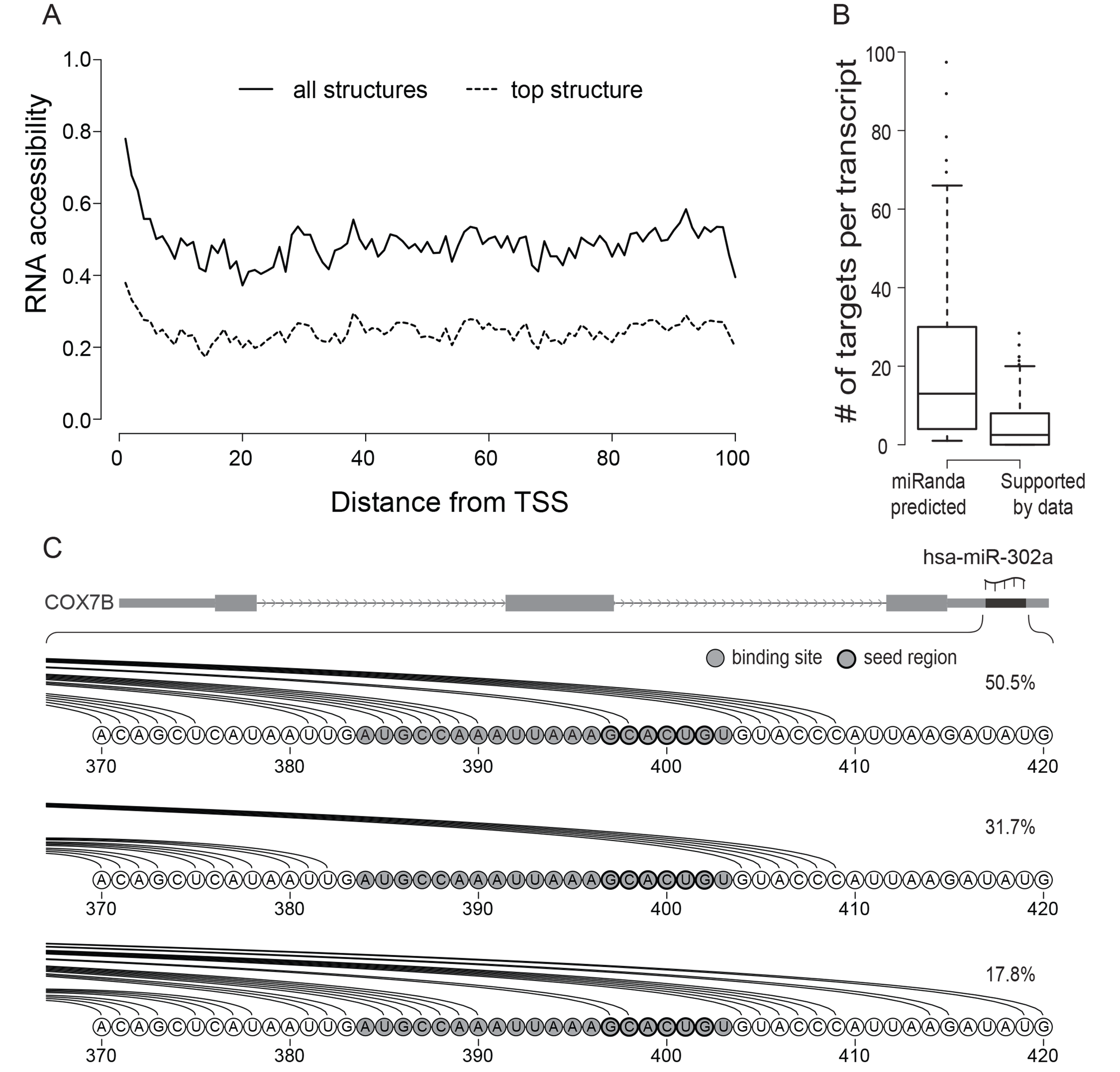
RNA accessibility analysis. (A) RNA accessibility at the 5’ end of mRNAs. Solid line represents the accessibility calculated from all RSQ quantified structures; dashed line represents accessibility calculated from the top structure of each transcript. (B) Boxplot of miRNA target site number for each transcript. The miRNA target site data obtained from http://www.microrna.org/; used human miRNA target site predictions with "good SVR scores and conserved miRNA" in August 2010 release. Target sites were filtered out if without RNA structural profiling data support. Distributions of target site number per transcript before and after filtering were shown f. (C) Accessibility of has-miR_302a target regions in transcript of COX7B gene. Quantified RNA conformation abundances weres shown. Nucleotides in gray Wang Page 29 Quantification of RNA structurome using next generation RNA structural profiling data circles are theoretically predicted hsa-miR-302a target sites. Paired bases are linked by arc lines.COX7B gene is found to be accessible for hsa-miR-302a to bind only in a less favored conformation.

## Discussion

We developed RSQ, a novel statistical model-based method for quantifying the RNA structurome using genome-wide RNA structural profiling data. The systematic and comprehensive analysis using the RSQ method outperformed previous analyses of RNA structural profiling data. RSQ shows higher tolerance to noise. The efficiency of the RNases varies depending on the versatile context of their digestion sites, thus substantial noise in local regions is inevitable. By calculating the maximum likelihood estimation (MLE) of the read number for each structure, the local noise was smoothed across the whole transcript, which led to more reliable results. Furthermore, RSQ makes use of experimental data to reduce the Boltzmann sampling space. Although the Boltzmann sampling method can nicely approximate the stationary distribution of conformation dynamics, for individual genes, the conformations theoretically predicted by Sfold (and/or other methods) may be inconsistent with RNA structural profiling data. By taking the top ranked signals (either single-strand or double-strand data) or mutual signals (both single-strand and double-strand data) as constraints into structure sampling/prediction, it guarantees the RSQ method to start with a more reliable structure sets. Finally and most importantly, RSQ provides a more meaningful interpretation of RNA structural profiling data based on the two layers of dynamics, the transcription dynamics and the RNA conformation dynamics, which is not provided by the existing analytic methods.

With emerging genome-wide RNA structural profiling data, the RSQ method makes it possible to understand RNA functions on the level of the conformation composition. Evidence has shown that variation in the RNA conformation composition can regulate gene expression levels by affecting the transcription efficiency and mRNA decay [43]. More interestingly, the RNA conformation composition may play a role in the regular function(s) of long RNA. Analyses of existing RNA structural profiling data in several human cell lines showed that the RNA conformation composition profile is relatively stable among cell lines, and its variation is not significantly correlated to gene expression variations. These findings also indicated a general regulation mechanism through which an RNA transcript can tune its function profile to some extent by changing its conformation composition, without affecting its transcription rate.

RNA structural profiling data are currently available for only a few cell lines. When the technologies are applied in more cell lines, especially tissue samples for various diseases, the quantified RNA structurome is expected to assist in deciphering disease-related RNA conformation composition variations. Genes without significant expression differences between samples might differ in RNA conformation composition. Moreover, armed with this genome-wide RNA structurome, the effects of single nucleotide variations (SNVs) on RNA conformation — dubbed ‘riboSNitches’ — can be surveyed [44]. Previously, a family trio study showed that riboSNitches constitute ~15% of all transcribed SNVs, which is far more than expected [10]. With the RSQ method, riboSNitches can be surveyed at a much finer resolution in consideration of gene isoforms and RNA conformation dynamics, which will allow for more accurate and extensive analyses of how SNVs change gene functions in states of health versus disease conditions.

A previous study reported that the structure-derived accessibility displays much higher correlation to translational efficiency than that derived from the raw sequencing signal [36]. RSQ further extends the structure-derived accessibility to that derived from the conformation dynamics. Instead of simply specifying the accessibility of a given region as “yes” or “no” from any single conformation for any single transcript transcribed from a gene locus, RSQ has the power to quantify the accessibility on the resolution of a single base pair based on the conformation dynamics for all the isoforms, which leads to more accurate evaluation of miRNA-mRNA or RBP-mRNA interactions and a more rational design of siRNAs in knockdown experiments.

## Acknowledgements

We thank William Su for a critical reading of the manuscript. This work was supported by the National Institutes of Health (HG0016960), National Natural Science Foundation of China (91019016), National Basic Research Program of China (2012CB316503), Cecil H. and Ida Green Distinguished Endowed Chair and the University of Texas at Dallas.

## References

1. Wan Y, Kertesz M, Spitale RC, Segal E, Chang HY: Understanding the transcriptome through RNA structure. Nat Rev Genet 2011, 12:641–655.

2. Mortimer SA, Kidwell MA, Doudna JA: Insights into RNA structure and function from genome-wide studies. Nat Rev Genet 2014, 15:469–479.

3. Mauger DM, Siegfried NA, Weeks KM: The genetic code as expressed through relationships between mRNA structure and protein function. FEBS Lett 2013, 587:1180–1188.

4. McManus CJ, Graveley BR: RNA structure and the mechanisms of alternative splicing. Curr Opin Genet Dev 2011, 21:373–379.

5. Warf MB, Berglund JA: Role of RNA structure in regulating pre-mRNA splicing. Trends Biochem Sci 2010, 35:169–178.

6. Martin KC, Ephrussi A: mRNA localization: gene expression in the spatial dimension. Cell 2009, 136:719–730.

7. Cruz JA, Westhof E: The dynamic landscapes of RNA architecture. Cell 2009, 136:604–609.

8. Garneau NL, Wilusz J, Wilusz CJ: The highways and byways of mRNA decay. Nat Rev Mol Cell Biol 2007, 8:113–126.

9. Kozak M: Regulation of translation via mRNA structure in prokaryotes and eukaryotes. Gene 2005, 361:13–37.

10. Wan Y, Qu K, Zhang QC, Flynn RA, Manor O, Ouyang Z, Zhang J, Spitale RC, Snyder MP, Segal E, Chang HY: Landscape and variation of RNA secondary structure across the human transcriptome. Nature 2014, 505:706–709.

11. Rouskin S, Zubradt M, Washietl S, Kellis M, Weissman JS: Genome-wide probing of RNA structure reveals active unfolding of mRNA structures in vivo. Nature 2014, 505:701–705.

12. Ding Y, Tang Y, Kwok CK, Zhang Y, Bevilacqua PC, Assmann SM: In vivo genome-wide profiling of RNA secondary structure reveals novel regulatory features. Nature 2014, 505:696–700.

13. Wan Y, Qu K, Ouyang Z, Kertesz M, Li J, Tibshirani R, Makino DL, Nutter RC,Segal E, Chang HY: Genome-wide measurement of RNA folding energies. Mol Cell 2012, 48:169–181.

14. Li F, Zheng Q, Vandivier LE, Willmann MR, Chen Y, Gregory BD: Regulatory impact of RNA secondary structure across the Arabidopsis transcriptome. Plant Cell 2012, 24:4346–4359.

15. Li F, Zheng Q, Ryvkin P, Dragomir I, Desai Y, Aiyer S, Valladares O, Yang JM, Bambina S, Sabin LR, et al: Global Analysis of RNA Secondary Structure in Two Metazoans. Cell Rep 2012, 1:69–82.

16. Lucks JB, Mortimer SA, Trapnell C, Luo S, Aviran S, Schroth GP, Pachter L, Doudna JA, Arkin AP: Multiplexed RNA structure characterization with selective 2’– hydroxyl acylation analyzed by primer extension sequencing (SHAPE-Seq). Proc Natl Acad Sci U S A 2011, 108:11063–11068.

17. Zheng Q, Ryvkin P, Li F, Dragomir I,Valladares O, Yang J, Cao K, Wang LS, Gregory BD: Genome-wide double-stranded RNA sequencing reveals the functional significance of base-paired RNAs in Arabidopsis. PLoS Genet 2010, 6:e1001141.

18. Underwood JG, Uzilov AV, Katzman S, Onodera CS, Mainzer JE, Mathews DH, Lowe TM, Salama SR, Haussler D: FragSeq: transcriptome-wide RNA structure probing using high-throughput sequencing. Nat Methods 2010, 7:995–1001.

19. Kertesz M, Wan Y, Mazor E, Rinn JL, Nutter RC, Chang HY, Segal E: Genome-wide measurement of RNA secondary structure in yeast. Nature 2010, 467: 103–107.

20. Spitale RC, Flynn RA, Zhang QC, Crisalli P, Lee B, Jung JW, Kuchelmeister HY, Batista PJ, Torre EA, Kool ET, Chang HY: Structural imprints in vivo decode RNA regulatory mechanisms. Nature 2015, 519:486–490.

21. Weeks KM: Advances in RNA structure analysis by chemical probing. Curr Opin Struct Biol 2010, 20:295–304.

22. Ehresmann C, Baudin F, Mougel M, Romby P, Ebel JP, Ehresmann B: Probing the structure of RNAs in solution. Nucleic Acids Res 1987, 15:9109–9128.

23. Ding Y, Chan CY, Lawrence CE: Sfold web server for statistical folding and rational design of nucleic acids. Nucleic Acids Res 2004, 32:W135–141.

24. Wu Y, Shi B, Ding X, Liu T, Hu X, Yip KY, Yang ZR, Mathews DH, Lu ZJ: Improved prediction of RNA secondary structure by integrating the free energy model with restraints derived from experimental probing data. Nucleic Acids Res 2015, 43: 7247–7259.

25. Bellaousov S, Reuter JS, Seetin MG, Mathews DH: RNAstructure: Web servers for RNA secondary structure prediction and analysis. Nucleic Acids Res 2013, 41: W471–474.

26. Black DL: Mechanisms of alternative pre-messenger RNA splicing. Annu Rev Biochem 2003, 72: 291–336.

27. Landry JR, Mager DL, Wilhelm BT: Complex controls: the role of alternative promoters in mammalian genomes. Trends in Genetics 2003, 19: 640–648.

28. Kowalczyk MS, Hughes JR, Garrick D, Lynch MD, Sharpe JA, Sloane-Stanley JA, McGowan SJ, De Gobbi M, Hosseini M, Vernimmen D, et al: Intragenic Enhancers Act as Alternative Promoters. Mol Cell 2012, 45:447–458.

29. Harrow J, Frankish A, Gonzalez JM, Tapanari E, Diekhans M, Kokocinski F, Aken BL, Barrell D, Zadissa A, Searle S, et al: GENCODE: the reference human genome annotation for The ENCODE Project. Genome Res 2012, 22:1760–1774.

30. Altuvia S, Kornitzer D, Teff D, Oppenheim AB: Alternative mRNA structures of the cm gene of bacteriophage lambda determine the rate of its translation initiation. J Mol Biol 1989, 210: 265–280

31. Weidner H, Yuan R, Crothers DM: Does 5S RNA function by a switch between two secondary structures? Nature 1977, 266: 193–194.

32. Jagadeeswaran P, Cherayil JD: A general model for the conformational switch in 5S RNA during protein synthesis. J Theor Biol 1980, 83: 369–375.

33. Ray PS, Jia J, Yao P, Majumder M, Hatzoglou M, Fox PL: A stress-responsive RNA switch regulates VEGFA expression. Nature 2009, 457: 915–919.

34. Kedde M, van Kouwenhove M, Zwart W, Oude Vrielink JA, Elkon R, Agami R: A Pumilio-induced RNA structure switch in p27–3’ UTR controls miR-221 and miR-222 accessibility. Nat Cell Biol 2010, 12: 1014–1020.

35. Sato K, Hamada M, Asai K, Mituyama T: CENTROIDFOLD: a web server for RNA secondary structure prediction. Nucleic Acids Res 2009, 37: W277–280.

36. Ouyang Z, Snyder MP, Chang HY: SeqFold: genome-scale reconstruction of RNA secondary structure integrating high-throughput sequencing data. Genome Res 2013, 277–387.

37. Li B, Dewey CN: RSEM: accurate transcript quantification from RNA-Seq data with or without a reference genome. BMC Bioinformatics 2011, 12: 323.

38. Li B, Ruotti V, Stewart RM, Thomson JA, Dewey CN: RNA-Seq gene expression estimation with read mapping uncertainty. Bioinformatics 2010, 26: 493–500.

39. Ding Y, Lawrence CE: A statistical sampling algorithm for RNA secondary structure prediction. Nucleic Acids Res 2003, 31: 7280–7301.

40. Enright AJ, John B, Gaul U, Tuschl T, Sander C, Marks DS: MicroRNA targets in Drosophila. Genome Biol 2003, 5:R1.

41. Betel D, Wilson M, Gabow A, Marks DS, Sander C: The microRNA.org resource: targets and expression. Nucleic Acids Res 2008, 36: D149–153.

42. Betel D, Koppal A, Agius P, Sander C, Leslie C: Comprehensive modeling of microRNA targets predicts functional non-conserved and non-canonical sites. Genome Biol 2010, 11: R90.

43. Vogel C, Marcotte EM: Insights into the regulation of protein abundance from proteomic and transcriptomic analyses. Nat Rev Genet 2012, 13: 227–232.

44. Lokody I: RNA: riboSNitches reveal heredity in RNA secondary structure. Nat Rev Genet 2014, 15: 219.

45. Ding Y, Chan CY, Lawrence CE: RNA secondary structure prediction by centroids in a Boltzmann weighted ensemble. RNA 2005, 11: 1157–1166.

